# The fitness consequences of coinfection and reassortment for segmented viruses depend upon viral genetic structure

**DOI:** 10.1101/2025.07.22.666171

**Authors:** Mireille Farjo, Christopher B. Brooke

## Abstract

Cellular coinfection between multiple virions is a common feature of viral infections. The collective virus-virus interactions enabled by these coinfections can influence the fitness of viral populations and give rise to novel infection phenotypes. Multi-strain coinfections allow viral resources to be shared between multiple individuals, and enable genetic combination and recombination between genotypes, potentially giving rise to hybrid progeny with enhanced fitness. However, coinfection can also impose fitness costs in certain situations. For example, resource sharing among viruses can lead to the persistence of low-fitness genotypes, and reassortment between different strains can lead to negative inter-segment epistasis when genes are poorly matched to one another. Thus, the fitness implications of cellular coinfection are poorly defined and likely context dependent. To investigate the specific conditions that lead to positive or negative fitness consequences for multi-strain coinfections, we formulated a model in which different genotypes of a three-segment virus replicate under varying degrees of inter-strain mixing. We observed that increased mixing had negative fitness consequences under a variety of scenarios, and that this effect was exacerbated with increasing genetic divergence between strains. Inter-strain mixing only enhanced viral fitness (a) when positive genetic dominance interactions were at play, and (b) under very specific conditions of selective pressure. We also observed that reassortment arising from mixing could generate hybrid genotypes with higher fitness than either parental virus, but that these outcomes were relatively rare. Overall, using a model segmented virus, we found that the heterologous coinfection was deleterious under most conditions, suggesting that it may be beneficial for many viruses to limit the extent of cellular coinfection.

## Introduction

Viral populations can encode a remarkable degree of genetic diversity^1–3^. Because of the high mutation rates observed across RNA viruses, even infections initiated by a single genome can generate distinct viral genotypes that may interact with each other during the course of infection^4–7^. Furthermore, coinfections between different strains can bring genetically divergent viruses into contact with each other, resulting in a spectrum of consequences that range from cooperation to competition.

During cellular coinfections, viruses may share gene products^8–12^ and even swap genetic material to generate hybrid progeny^7,13–16^. Genetic exchange can occur via copy-choice recombination, in which the viral polymerase switches templates between different strands of nucleic acids mid-replication, resulting in a hybrid molecule^17^. Alternatively, for viruses with segmented genomes, genetic exchange can also be achieved through reassortment, which occurs when gene segments from different strains are co-packaged in the same viral particle. This process can be highly efficient, and extensive reassortment between segmented viruses has been observed in the wild^18–21^.

Previous studies have suggested multiple possible benefits of cellular coinfection. One such advantage is facilitating functional innovation through cooperation between different strains. For example, interactions between distinct genotypes during infection can support novel collective viral traits like cell-to-cell spread^22–25^ and expanded tropism^26^. Coinfection also enables genetic exchange, which can lead to increased adaptability in cases where recombination and reassortment allow viruses to reorganize alleles into more optimal configurations. For example, the experimental generation of hybrid H3N2/H5N1 influenza viruses revealed that multiple reassortant strains exhibit higher replicative capacity than either parental virus^27^, and studies on poliovirus have demonstrated how recombination can increase the rate of adaptation during tissue culture infection^28^. Phylogenetic analyses have also shown that flu lineages derived from reassortant strains persist longer than their non-reassortant counterparts^29^, and similar approaches have been used to show that higher rates of recombination correlate with greater adaptability in coronaviruses^30^. One of the major manifestations of genetic exchange-derived adaptability is the rapid introduction of new antigens into an immunologically naïve population, a feature illustrated by the fact that every influenza pandemic of the past century has been driven by a reassortant virus^31^.

Genetic exchange could also promote adaptability by allowing viruses to maintain a high mutation rate. Mutational diversity can be associated with increased adaptive capacity^32,33^, but since the majority of mutations are deleterious, high mutation rates can also decrease overall fitness and inhibit the emergence of beneficial alleles^34,35^. Recombination and reassortment can offset the costs of a high mutation rate by allowing low-fitness alleles to be quickly replaced by alternatives and swept out of the population^7,36^. Achieving an optimal balance between mutation rate and recombination rate can thereby allow a virus to maximize its fitness during infection.

Despite the potential benefits of cooperative viral interactions and genetic exchange, many viruses encode mechanisms to limit the extent of cellular coinfection. These mechanisms come into play after one virus initiates an infection, causing a downstream effect that blocks subsequent infection of the same cell. This process, known as superinfection exclusion, can be observed across distantly related viruses, including influenza^37–39^, SARS-CoV-2^40^, West Nile virus^41,42^, vaccinia virus^43^, turnip crinkle virus^44^, phage λ^45^, and phage T4^46^. Superinfection exclusion has also been shown to impact the tissue-scale patterning of infection within a host, with viral populations segregating into “infection islands” that have a limited capacity to inter-mix^38,47^.

The extent to which superinfection exclusion is observed across a diverse array of virus families raises the question of whether inhibiting coinfection may be adaptive for the virus. Although coinfection (and the consequent enabling of genetic exchange) can give rise to successful viral lineages, it is possible that these high-fitness recombination events are relatively rare. Sequence analysis of seasonal H3N2 influenza populations has shown that reassortment is observed less frequently than would be expected under a null model, suggesting that a substantial proportion of reassortment events may yield progeny that are less fit than their parents^48^.

There are multiple mechanisms by which coinfection can promote the generation and/or maintenance of low-fitness genotypes. For instance, negative epistatic interactions between gene segments could lead to the generation of low-fitness reassortant genomes. Inter-segment epistasis can occur when gene products need to physically interact with each other — for example, the three genes that comprise the influenza replicase form a multimeric protein complex, and genetic mismatches between segments can disrupt the necessary configuration^15,49^. Alternatively, viral genes may encode opposing functions, which must be balanced to maximize the fitness of the virus as a whole^50–53^. Under these circumstances, reassortment may be more likely to generate a faulty virus than a high-fitness or neutral one.

Furthermore, coinfection can result in the complementation of deleterious mutations, allowing them to persist longer in a population^54–56^. Genetic complementation can also support populations of non-standard viral genomes (such as defective viral genomes (DVGs) and/or deletion-containing viral genomes (DelVGs)), which are unable to replicate independently of a helper virus and, in some cases, have been shown to interfere with infection^57–60^. These negative consequences of complementation may outweigh the positive effects of cooperation between coinfecting genotypes.

We aimed to explore the potentially contrasting fitness implications of cellular coinfection using a modeling approach. We simulated infections with a model three-segment virus while modeling varying degrees of inter-strain interactions under different conditions. We found that mixing between viral populations tended to result in negative fitness outcomes, but that immune pressure directed at antigenic genes could, in some cases, promote more positive coinfection outcomes. We also observed that coinfection dampened the correlation between an individual’s fitness and its genetic representation in the following generation, driven by a relaxation of selection on individual genotypes. Our results reveal multiple factors that shape the fitness effects of multi-strain coinfections and suggest an adaptive logic for the restriction of virus-virus interactions during infection.

## Results

### Model overview

We aimed to investigate the implications of heterotypic coinfection on viral fitness by simulating the effect of a range of virus-virus coinfection rates on replication and packaging. To this end, we designed a model virus expressing three gene segments (A, B, and C), which are assigned intrinsic activity values that, together, determine the overall genotypic fitness. This genome structure allowed us to explore different features of segmented viruses such as influenza viruses and hantaviruses within our model. Each simulated coinfection is initiated by a pair of parental genotypes, which encounter each other according to a predefined mixing rate. We observed the effect of the mixing rate on overall viral replication, as well as on the fitness of the resulting genomes.

### Defining genotypes

We assigned each parental genotype a per-gene activity value *(x)* for each of the segments A, B, and C. These values were assigned randomly from a distribution **(Figure 1A)** adapted from a previously reported mutational fitness effects distribution for the HA gene of influenza A virus, generated through deep mutational scanning (DMS)^53^. To adapt the DMS-derived distribution for our purposes, we replaced every score ≤ 0 with a value of 0.00001, but the dataset was otherwise unmodified.

**Figure 1.**
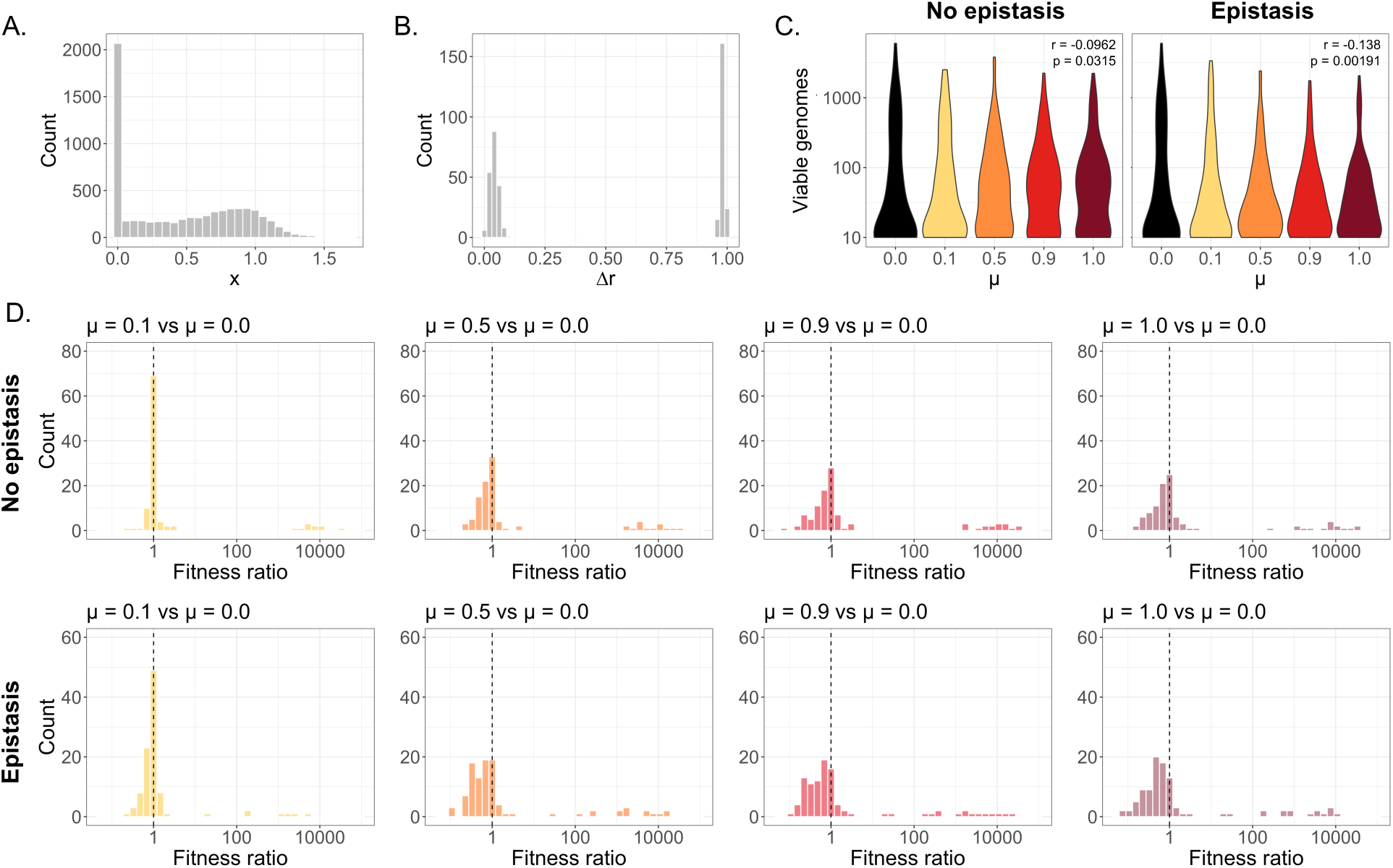
Replication under non-epistatic and epistatic scenarios. **(A)** Distribution of the per-segment activity metric x. **(B)** Distribution of the phylogenetic distance metric Δr. **(C)** Total viable genomes under varying values of μ, where distributions represent stochastic simulation outputs for 100 coinfecting genotype pairs. Insets show *ρ* and p-values, derived from a Spearman’s rank correlation test. **(D)** Ratio of weighted offspring-population fitness scores between mixed and unmixed conditions, calculated for individual coinfections. Dashed line represents a fitness ratio of 1.0, at which the fitness of the unmixed and the fully mixed populations are equivalent; values > 1.0 represent simulations in which overall fitness was higher for mixed populations relative to unmixed populations.

To model the effect of strain genetic divergence on coinfection (which affects the extent of inter-segment epistasis), each coinfecting pair was also assigned a dimensionless value (Δ*r*), representing the phylogenetic distance between the strains **(Figure 1B)**. To determine the distribution of Δ*r* values, we first downloaded all PB2 gene sequences of human-sampled H1N1 and H3N2 originating from the state of Texas that were submitted by the CDC to GISAID during the 2024-2025 flu season (from 11-01-2024 to 05-01-2025). We then calculated Nei’s nucleotide diversity within H1N1 or H3N2 sequences, and between H1N1-H3N2 pairs, and generated equally sized normal distributions based on the mean and standard deviations of the nucleotide diversity values for each coinfection type (H1N1-H1N1, H3N2-H3N2, H1N1-H3N2, and H3N2-H1N1). These values were then normalized so that the final distribution of *Δr* ranged from 0 to 1. We defined the post-epistasis fitness of each gene as:

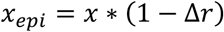

Thus, when *Δr* = 0, epistasis has no effect on fitness, and when *Δr* = 1, epistasis renders the gene nonfunctional. This range of impacts reflects previous observations that genetic mismatch between viral gene segments can lead to nonviability^61^.

### Simulating replication and packaging

Coinfections were initialized with 5 founder genomes derived from each parental genotype (genotypes i and j). Every infecting genome was complete, containing an A, B, and C segment. Coinfecting genomes were sorted into two viral subpopulations, with the distribution of genotypes i and j across the subpopulations defined by a mixing rate (*μ*). We established a set of *μ* values ranging from 0-1, where *μ* = 0 signifies complete segregation of parental genotypes and *μ* = 1 indicates complete mixing. The number of founding genomes (*f*) of genotype i that are sorted into subpopulations i and j is defined by:

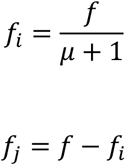

Therefore, at *μ* = 0, coinfecting populations are completely segregated into separate subpopulations such that a virus of genotype i has no chance of encountering a virus of genotype j. At *μ* = 1, the populations are completely mixed, and the chance of a virus encountering its own genotype versus the coinfecting genotype depends only on their relative population frequencies.

We first considered a scenario in which viruses do not experience inter-segment epistasis, and the fitness (*w*) of each allele is equal to its intrinsic activity (*x*). Because we expect intracellular resources to be shared, we calculated the per-gene fitness as the weighted average of the activities of each allele in the subpopulation. We also assumed that every gene segment is necessary for productive infection, such that the lowest-fitness gene serves as the limiting factor in defining the overall subpopulation fitness. We therefore defined the overall subpopulation fitness as:

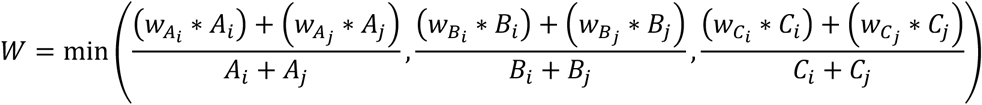

Where *A_i_*, *A_j_*, *B_i_*, *B_j_*, *C_i_*, and *C_j_* are equal to the total number of the alleles in the population at a given time point.

When *μ* < 1, the initial value of *W* will differ between subpopulations because of differences in the frequencies of each parental genotype. However, when *μ* = 1, genotypes i and j are distributed equally across both cells, and the initial value of *W* will be the same for each subpopulation.

We then defined per-segment replication propensities as:

**Table 1.**
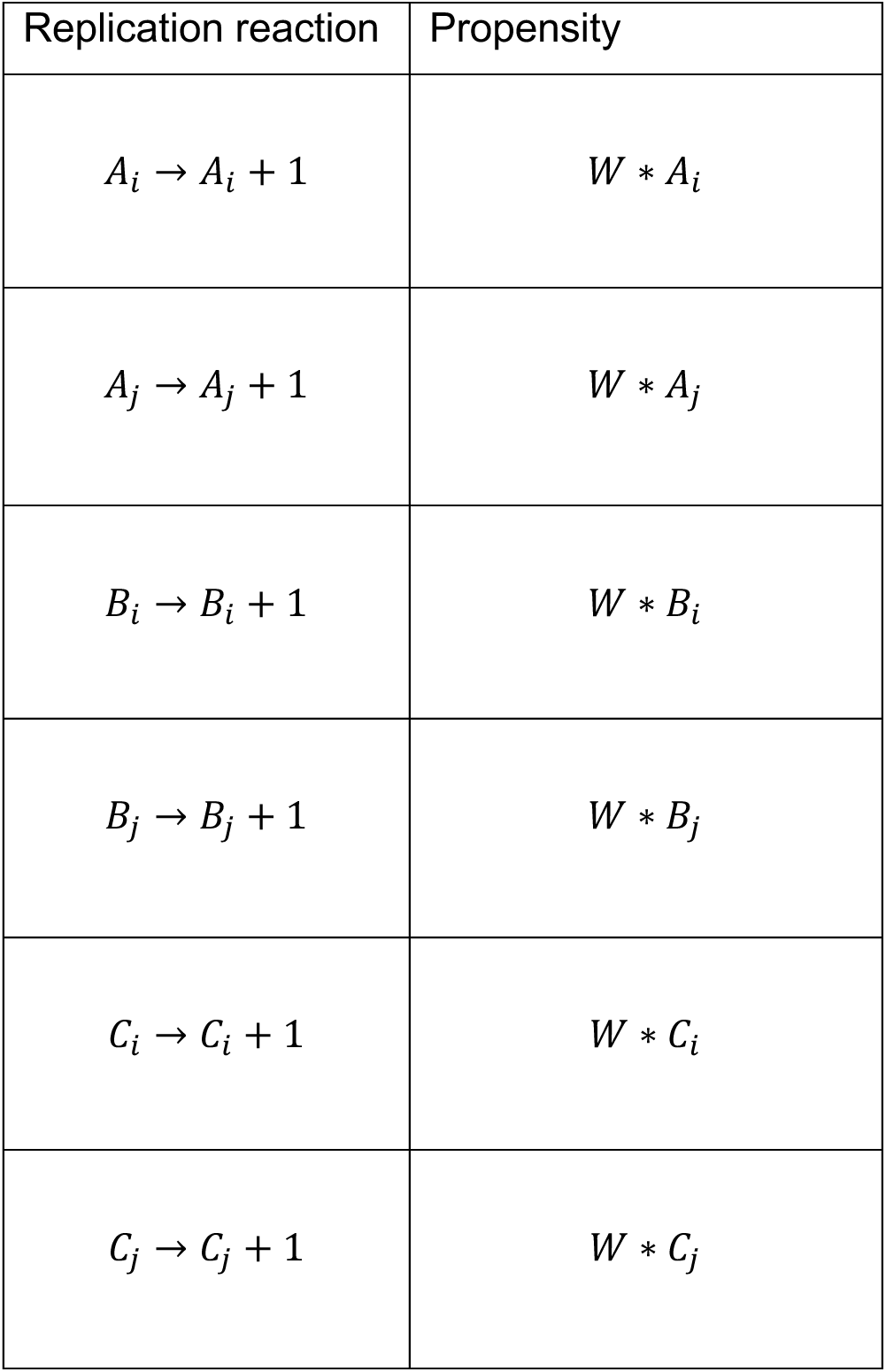
Per-segment replication propensities.

Replication propensities were also contingent on the current count of a given segment (along with the overall fitness) because we expect a gene’s likelihood of interaction with replication/translation machinery to depend in part on its frequency within the cell.

We calculated per-segment replication propensities for both subpopulations, based on the frequency of each gene segment within the given cell. We then modeled viral replication for both subpopulations using a Gillespie stochastic simulation algorithm^62^. After each replication event, segment counts were updated accordingly, along with the corresponding per-segment and per-subpopulation fitness values. Simulations were performed separately for each subpopulation, so that separate infected cells did not directly compete with one another.

After simulating a single cycle of replication for each coinfecting genotype pair, we calculated a “viable genomes” metric, defined as the maximum number of complete genomes (containing a single copy of segments A, B, and C) that could be constructed from each subpopulation’s post-replication gene pool. We then pooled the viable genomes from both subpopulations to yield a total viable genomes count, calculated under varying values of *μ* **(Figure 1C)**.

In the absence of inter-segment epistasis, we observed a slight negative correlation between mixing rate and replication, possibly caused by differences in the distribution of viruses of varying fitness values across subpopulations. In our model, the replicative capacity of a given subpopulation depends on the fitness contributions of each constituent gene. When replication occurs exponentially, concentrating high-fitness genes within the same subpopulation results in greater overall replication than when fitness benefits are averaged out across multiple subpopulations. Therefore, dispersing a high-fitness virus and a low-fitness virus equally across two compartments (as when *μ* = 1) results in less replication overall than if each virus is restricted to its own spatial arena. Our results suggest that segment incompatibility is not the only factor that can lead to decreased viral output during mixed coinfections, and that negative replication consequences may occur when a high-fitness strain sees its replication capacity diminished through intracellular interactions with a low-fitness coinfection partner.

However, despite the negative relationship between mixing rate and replication, and although the mean fitness of the mu = 1 population was lower than that of the mu = 0 population (mean = 145.26 vs. mean = 294.08), we observed that the median fitness of the fully mixed population was higher than that of the unmixed population (median = 49.5 vs. median = 21). This relationship reflects a role for mixing in rescuing infections that might otherwise be non-productive.

We next considered a scenario in which the fitness of our model virus was constrained by inter-segment epistasis. To this end, we defined segments A and B as members of an epistatic pair and updated our per-segment fitness values so that the fitnesses of segments A and B were dependent on both their intrinsic activities (*x*) and the likelihood of encountering a homotypic partner versus a heterotypic partner. Under each encounter scenario, the encounter likelihood depends on the population frequency (*p*) of the partner, and the fitnesses of A and B are modified by the corresponding differences in cladal position (Δ*r*) of the interacting partners, such that a greater difference in *r* results in a lower fitness. We thus define the fitness for each segment (A, B, and C) of each genotype (i and j) as:

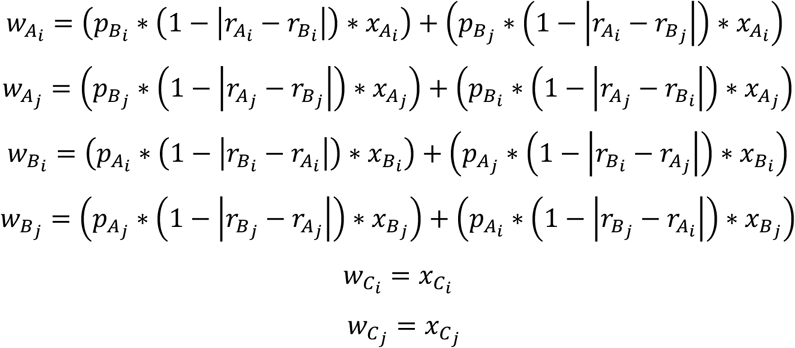

As previously, the overall subpopulation fitness was limited by the lowest-fitness gene of the three genomic segments.

Under conditions of inter-segment epistasis, we observed a greater negative impact of mixing on genome replication than under the unconstrained condition **(Figure 1C)**. Because we assumed that the genotypes of each parental virus were internally well-related, inter-strain mixing tended to generate A/B gene combinations that were (at best) about equally as related as the parental pairs, and that were frequently only distantly related. Due to negative epistasis, interactions between distantly related genes resulted in decreased replication outputs as mixing rates increased. Inter-strain epistasis therefore acted to exacerbate the negative impacts of heterologous mixing.

After generating post-replication pools of gene segments for each subpopulation, we also simulated viral packaging by assigning the segments to new genome sets. We assumed that packaging would occur with equal efficiency regardless of the identities of the co-packaged segments, and so the likelihood of each possible genomic combination depended only on the relative frequency of each constituent allele within the subpopulation. (See Methods.) We allowed for the packaging of every possible reassortant genotype, but we did not generate incomplete genomes missing one or more segments.

After simulating the packaging of gene segments into new combinations, we calculated the fitness of the resulting genomes, defined as the minimum per-gene fitness (*w*) of the three packaged segments (and considering the genetic distance between segments A and B under the epistatic scenario). We then calculated a weighted population fitness metric, such that the overall population fitness was defined by the relative proportion of each constituent genotype and its respective fitness value.

While inter-strain mixing had an overall negative impact on replication, we wanted to determine whether there were specific cases in which mixing led to the generation of higher-fitness progeny. To this end, we calculated the ratio of post-packaging fitness scores between mixed and unmixed populations for each simulated coinfection. As expected, post-replication population fitness tended to decrease under higher levels of mixing, and most individual coinfections saw lower fitnesses when any of the tested mixing rates were in place **(Figure 1D)**. These findings reflect the capacity of coinfection to support the propagation of low-fitness genotypes through functional complementation, leading to the persistence of these strains in the population. However, gene-swapping between strains can also support the generation of high-fitness genotypes by replacing nonfunctional alleles with viable alternatives. This outcome was slightly less common under the epistatic scenario, given the genetic constraints imposed by inter-segment interactions, but we still observed multiple individual cases where reassortment allowed for a (sometimes drastically) higher-fitness offspring generation **(Figure 1D)**.

Because of our finding that reassortment could lead to higher population fitness in certain cases, we then calculated maximum genotypic fitness values for each population **(Figure 2)**. We observed that only a low level of mixing was necessary to reliably generate these high-fitness genomes, both in the presence and absence of epistasis, and that maximum fitness distributions were essentially unchanged within a *μ* range of 0.5-1.0. Altogether, these data indicate that mixing is generally detrimental for average population fitness, but can promote the generation of novel, high-fitness genotypes not accessible in the absence of mixing.

**Figure 2.**
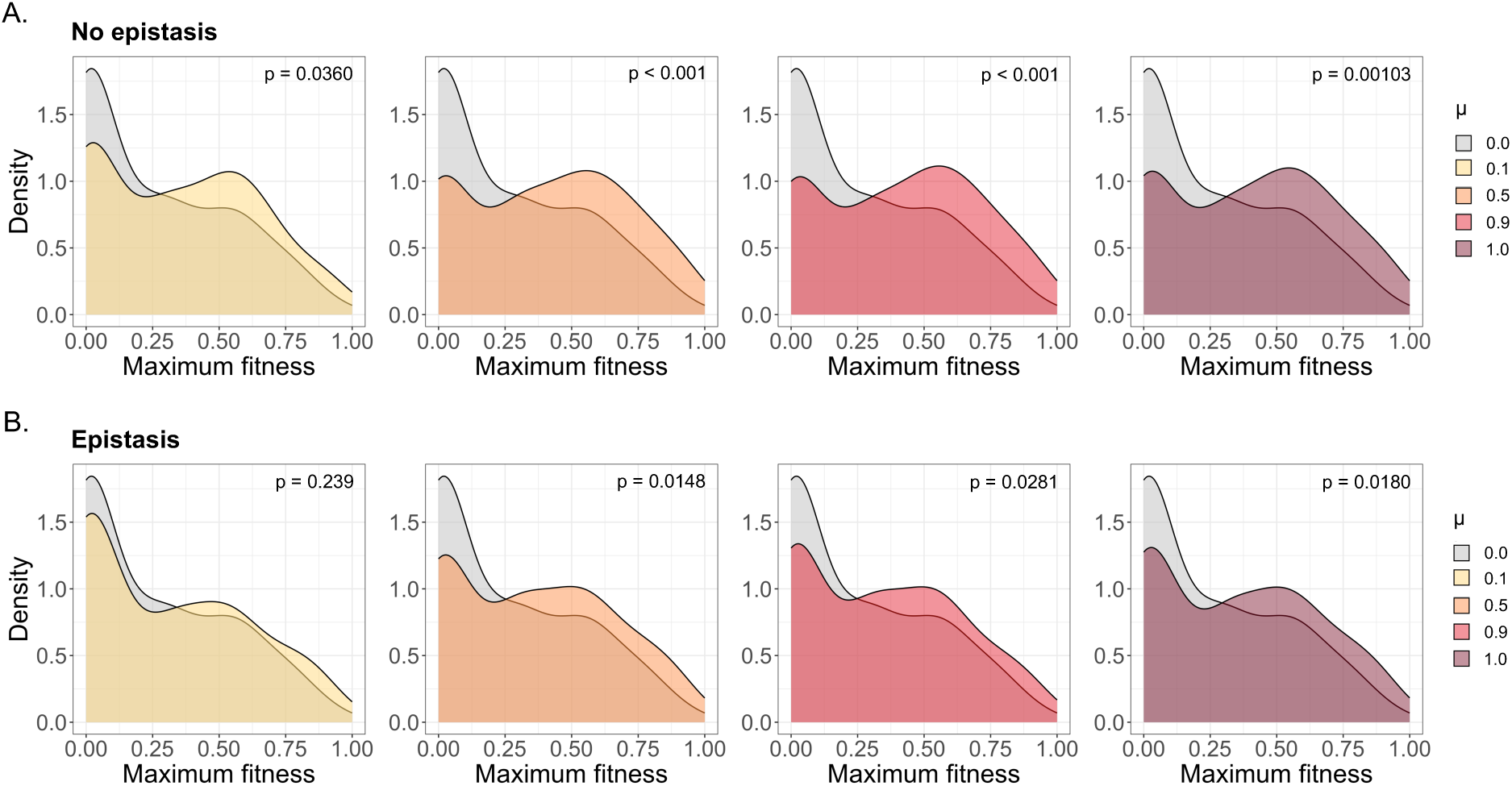
Maximum population fitness in non-epistatic and epistatic scenarios. **(A)** Distributions of the highest individual fitness values observed in each of the 100 coinfection simulations, under varying degrees of mixing and no inter-segment epistasis. P-values are derived from Mann-Whitney U tests comparing the fitness distributions of each given population. **(B)** Maximum fitness distributions for simulations under inter-segment epistasis.

### Effects of mixing under alternative genetic dominance structures

We performed our initial simulations under the assumption that different alleles within a subpopulation would contribute equally to the overall functionality of the gene in question. However, alleles may also interact through more complex dominance dynamics. For example, defective polymerase proteins translated from deletion-containing viral genomes can inhibit viral replication in a dominant-negative fashion^63^. Alternatively, a high-fitness viral allele that produces a public good protein (*e.g.* a potent ß immune antagonist) in abundance may have a dominant-positive effect on the overall population, compensating for any non-producer alleles without incurring a shortage of the protein in question.

In our initial simulations, we also assumed that the overall fitness of a viral genome would be entirely limited by its least-fit gene: however, under some circumstances, one high-fitness gene may partially compensate for a low-fitness allele on the same genome. For example, a fast-replicating virus may outpace the response of the innate immune system, rendering viral immune antagonism less crucial for productive infection^64^. We therefore wanted to explore the impact of coinfection under different scenarios of genetic interaction between alleles.

To investigate the replication impacts of heterologous coinfection under a positive-dominance circumstance, we modeled a scenario in which a pool of high-fitness alleles can cancel out the impact of an equally sized pool of their lower-fitness counterparts. Under this scenario, we defined per-gene fitness as:

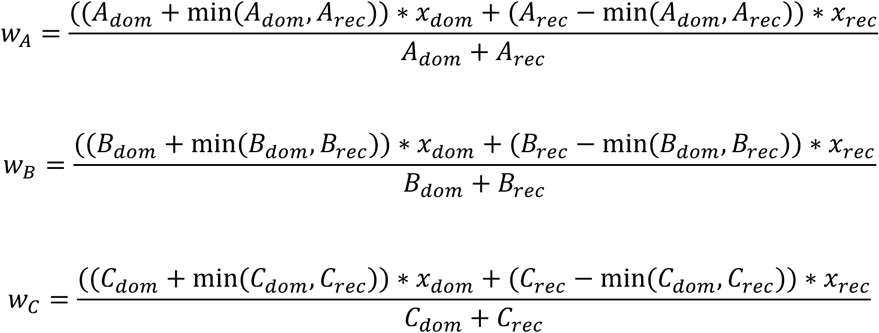

Where A_dom_, A_rec_, B_dom_, B_rec_, C_dom_ and C_rec_ represent the total copy number of the dominant (higher-fitness) and recessive (lower-fitness) alleles for each gene. Contrary to our previous observations, under this scenario, we observed that genetic mixing had a positive impact on overall replication **(Figure 3A)**. Under fully mixed conditions, both subpopulations contain roughly the same number of high-fitness (A_dom_) and low-fitness (A_rec_) alleles, and because each dominant, high-fitness allele has the capacity to overwrite the negative fitness impacts of a corresponding recessive allele, gene fitness can be maximized across both subpopulations at once. However, positive dominance also promotes the rescue of low-fitness alleles through coinfection with other strains — therefore, the fitnesses of individual genotypes packaged post-replication were still generally lower under increased mixing **(Figure 3B)**.

**Figure 3.**
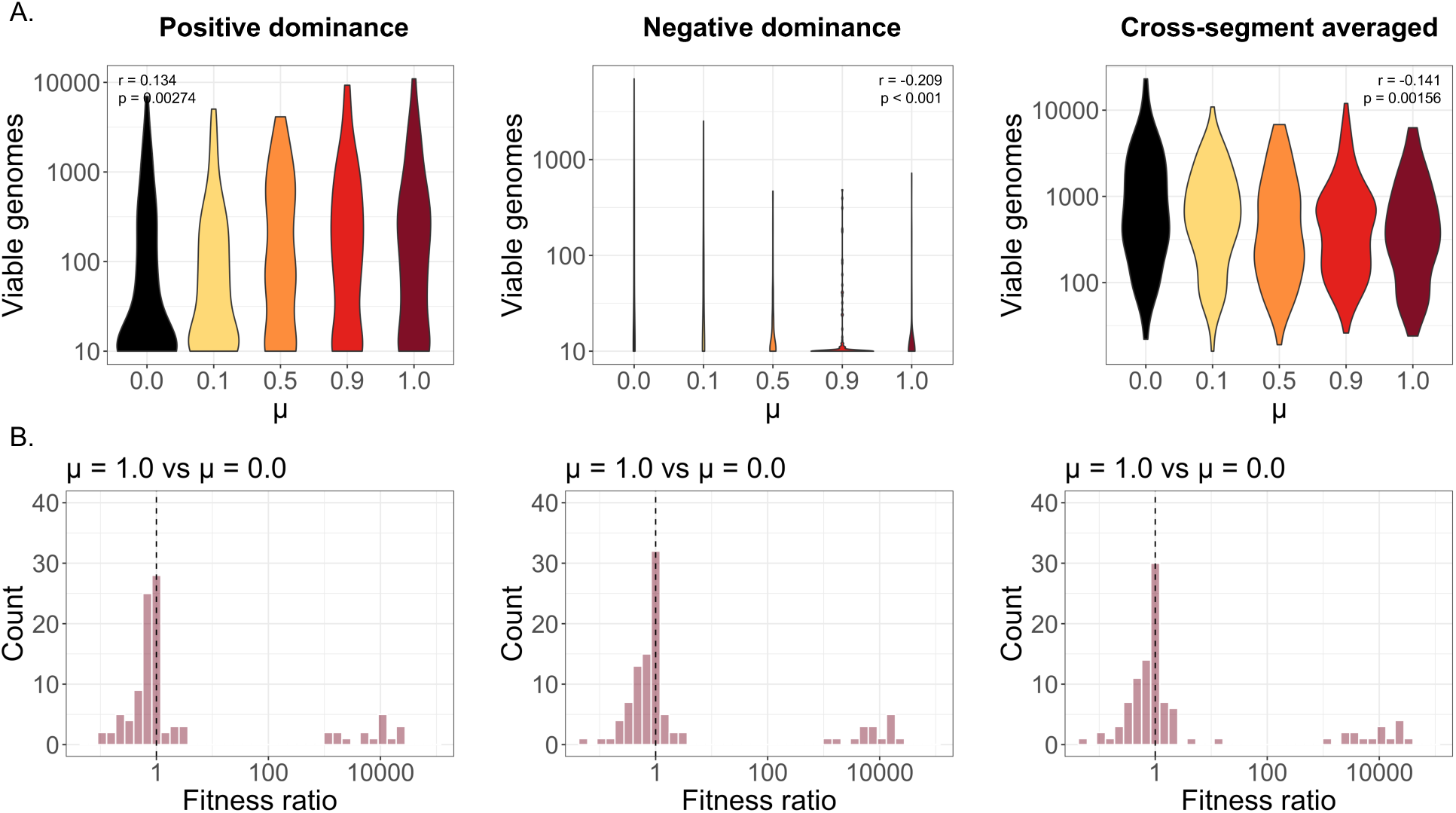
Replication and weighted population fitness shown for different methods of calculating replication propensities. **(A)** Viable genome counts for increasing values of μ under positive genetic dominance, negative genetic dominance, and cross-segment averaged fitness. **(B)** Ratio of weighted offspring-population fitness scores between fully mixed (μ = 1.0) and unmixed (μ = 0.0) conditions. Dashed line represents a fitness ratio of 1.0.

We then repeated the same simulation for the negative-dominance scenario, this time designating the dominant allele as the lower-fitness representative. As expected, we saw that negative dominance dynamics resulted in a pronounced negative relationship between mixing rate and replication, because mixing facilitates the spread of low-fitness alleles that can poison the replication of functional variants **(Figure 3A)**.

Finally, we considered a scenario in which the overall subpopulation fitness is not limited by the lowest fitness gene but is instead defined by the fitness of all three gene segments. Under this condition, we calculated overall fitness as the weighted mean of all alleles in the given subpopulation:

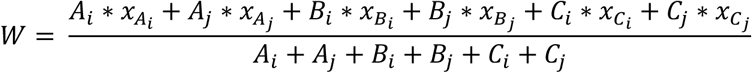

After repeating our simulations using the updated fitness calculations, we observed that higher mixing rates significantly decreased the overall level of viral replication **(Figure 3A)**. Additionally, the negative correlation between mixing and replication was greater than the one observed in the previously simulated no-epistasis condition, under which overall fitness depended only on the lowest-fitness gene of the three segments **(Figure 1C)**. One of the immediate benefits of gene flow is that it allows for the complementation of low-fitness alleles by more functional ones. However, when overall fitness depends equally on all three gene segments, there exist increased opportunities for high-fitness genes to compensate for a lack of activity in lower-fitness genes, even in the absence of genetic mixing. Therefore, the role of mixing in genetic complementation is diminished, and its negative impacts are balanced to a lesser degree.

### Coinfections under immune pressure

Because one of the potential benefits of reassortment is the rapid spread of resistance alleles (*e.g.* antigenic variants or drug resistance alleles), we decided to model a condition in which certain genes encode immunodominant antigens subject to immune pressure. For these antigenic genes, we defined a new activity distribution where the bulk of variants were nonviable (x = 0.00001; models antigenic sequences effectively recognized by host immune memory), and the activity scores of resistant mutants were assigned from the pool of viable variants (x > 0.00001) originating from the DMS dataset^53^ shown in Figure 1A **(Figure 4A)**. We modeled an initial population resistance level of 15%, and applied the resulting distribution to three genomic schemes: one (A = B = C) in which none of the three genes act as antigens under immune pressure, another (AB < C) in which segments A and B are antigens, and a final scheme (C < AB) in which segment C is an antigen. We again generated 100 pairs of coinfecting genotypes for each schematic, and simulated replication and packaging using a Gillespie algorithm.

**Figure 4.**
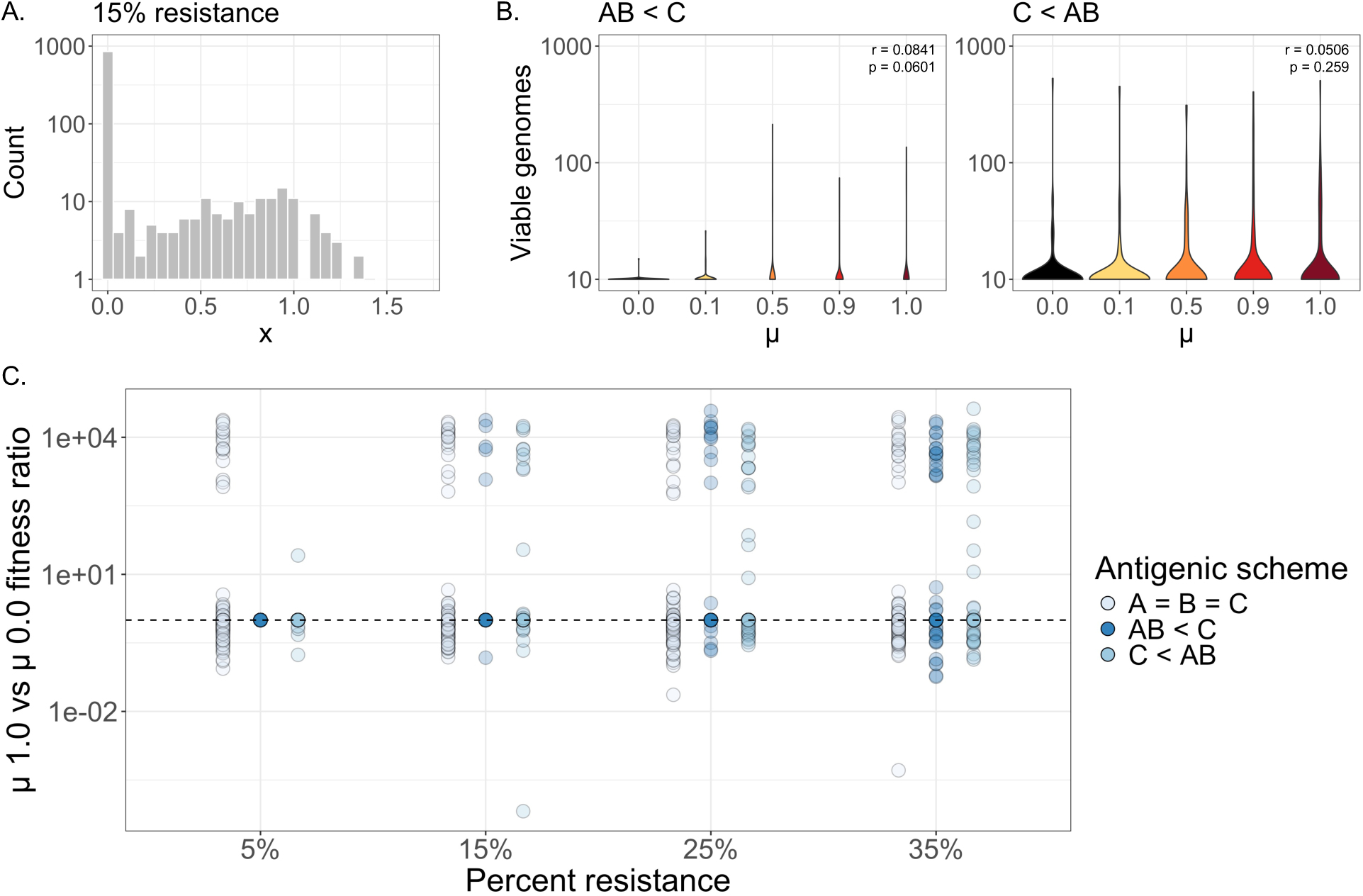
The impact of immune pressure on replication and fitness. **(A)** Distribution of gene activity (x) values under an immune-pressure model, where 15% of variants encode some degree of resistance. **(B)** Total viable genomes under different mixing rates (μ), for AB < C and C < AB antigenic schemes. Insets show *ρ* and p-values, derived from a Spearman’s rank correlation test. **(C)** Ratio of weighted population fitness scores between fully mixed (μ = 1.0) and unmixed (μ = 0.0) populations, shown in the absence of immune pressure (A = B = C) or under different immunity schemes (AB < C or C < AB) under varying levels of population resistance. Dashed line represents a fitness ratio of 1.0.

After calculating total viable genomes for the two antigenic conditions, we observed that increasing *μ* values did not have a significant effect on viable genome production for either the AB < C or the C < AB scheme **(Figure 4B)**. We next considered the possibility that the benefits of reassortment under immune pressure may depend on the frequency of resistance alleles pre-existing within the population, in addition to the number of antigens. We repeated our simulations in populations encoding levels of resistance ranging from 5-35% and calculated viable genome metrics under each resistance condition **(Supplemental Figure 1).** We observed that the 15% resistance model **(Figure 4B)** was the only scenario in which populations under the AB < C antigenic scheme saw a positive replication effect as a result of mixing, and that C < AB populations were overall unimpacted by mixing. We also calculated fitness ratios between fully mixed (*μ* = 1.0) and unmixed (*μ* = 0.0) populations at different resistance levels **(Figure 4C)**, and determined that mixing in the immunity scenarios (AB < C) and (C < AB) generally had a similar impact on population fitness as in the non-immunity scenario (A = B = C), regardless of the degree of immune resistance present in a population.

Finally, we considered the possibility that epistatic interactions between antigenic genes (as observed for influenza A virus HA and NA^52,65,66^) may also constrain the benefits of genetic mixing in populations under immune pressure. We repeated our simulations, accounting for epistasis between segments A and B as detailed previously, and calculated replication rates and fitness values for populations under both antigenic schemes **(Supplemental Figure 2).** We observed similar results as under the non-epistatic scenario, with only the AB < C population at 15% resistance seeing (minor) increases in replication under higher values of *μ*. Altogether, these findings suggest that gene flow may only facilitate immune escape under highly specific conditions depending on the frequency of resistance alleles in the population and the antigenic structure of the virus.

### Generational changes in allelic diversity

Reassortment and inter-strain mixing can influence the overall amount of allelic diversity within populations. To assess how varying levels of mixing affected population composition between parental and offspring generations, we examined the relationship between the relative fitness of parental genotypes originating from i (plotted as the fitness difference between parental genotypes i and j) and the generational change in frequency of those genotypes **(Figure 5)**.

**Figure 5.**
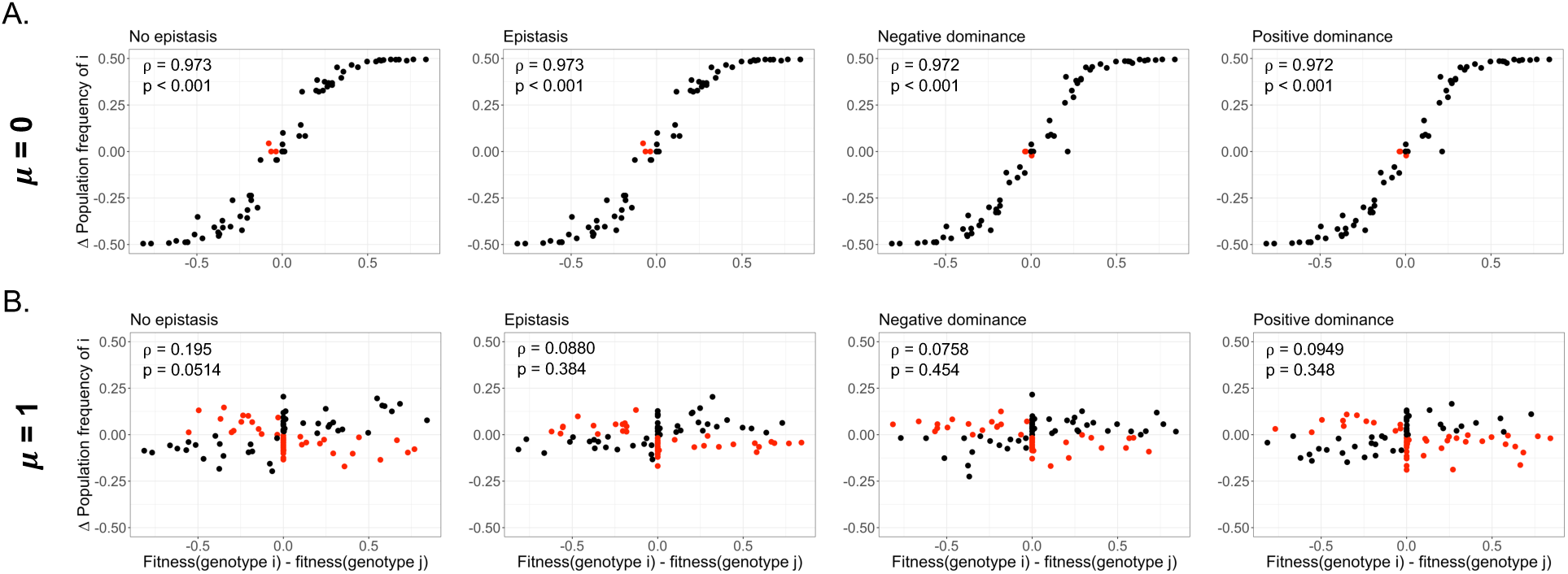
Genotypic fitness difference between genotypes i and j versus the post-replication frequency change of i alleles. Relationships are plotted for unmixed **(A)** or fully mixed **(B)** conditions. Red points indicate instances where alleles originating from a more fit genotype decrease in frequency, or vice-versa. Insets show *ρ* and p-values, derived from a Spearman’s rank correlation test.

Post-replication gene frequencies were calculated from a gene pool containing all packaged genomes, including reassortant genotypes. As the initial population frequency of each parental genotype (and its constituent alleles) was 0.5, a change in frequency of ± 0.5 indicated that the given allele had either reached fixation or gone extinct.

As expected, under unmixed conditions, there was a strong positive correlation between a genotype’s fitness and its degree of representation in the subsequent generation **(Figure 5A)**. However, when populations were fully mixed, this correlation between fitness and post-replication population frequency disappeared **(Figure 5B)**. This indicates that, when mixing levels were high, deleterious alleles were able to persist for longer in the gene pool, as their negative fitness impacts could be partially balanced by high-fitness coinfection partners. Post-packaging, reassortment may allow alleles like these to swap into a genomic context where their effects would be more favorable, or to persist by hitchhiking on the backbone of higher-fitness genotypes. These results indicate how viral mixing not only contributes to population diversity through the generation of new genotypes but also serves to maintain standing genetic diversity.

### Post-transmission replication and fitness

Because of the differences in population composition that we observed between mixed and unmixed infections, we wanted to explore the consequences of these differences for future infections. To simulate a second round of infection, we selected two viral genotypes from each of the simulated mixed (μ = 1.0), partially mixed (μ = 0.1), and unmixed (μ = 0.0) post-replication pools to initiate a new round of coinfections. This small number of founder genomes was chosen to model the narrow transmission bottlenecks that have been measured for IAV and SARS-CoV-2^67–69^. Because we selected genotypes based solely on their population frequency, the two selected viruses could be either homotypic or heterotypic. We then simulated a second round of replication and packaging for each coinfection pair, under mixed or unmixed conditions.

After calculating total viable genomes, we saw that there was no significant difference between the replication capacities of viruses drawn from the μ = 0.0, μ = 0.1, or μ = 1.0 under either unmixed or mixed conditions **(Figure 6A)**, suggesting that the bottleneck between the first and second generations mitigated replication defects by removing certain deleterious alleles from the population. We also observed that, under unmixed conditions, viruses drawn from the μ = 0.1 pool replicated to slightly higher levels than those drawn from the μ = 0.0 pool (median = 34.0 vs. median = 25.0; mean = 533.33 vs. mean = 518.93).

**Figure 6.**
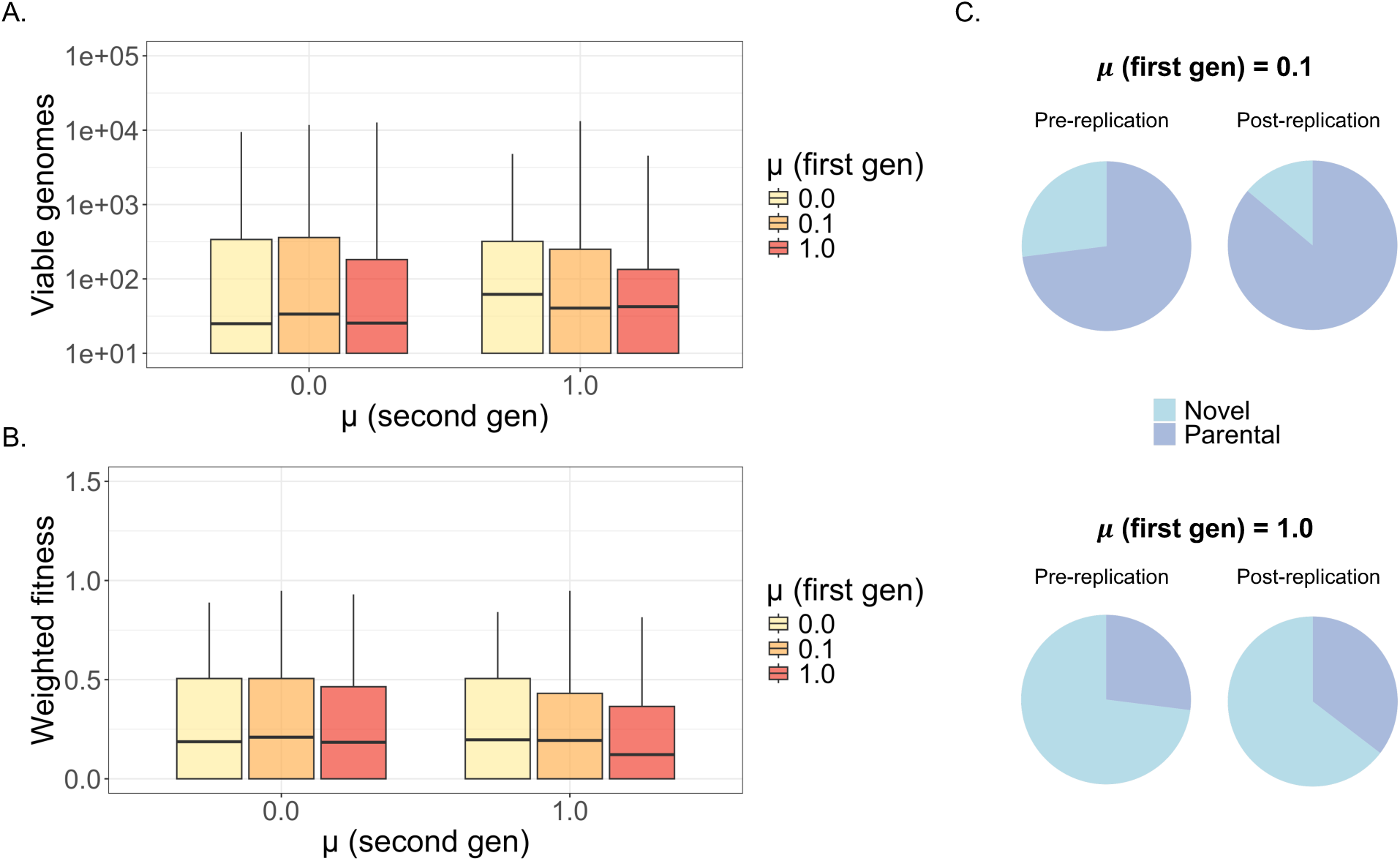
The impact of mixing in a second generation of replication. **(A)** Total viable genomes from coinfections initiated by viruses drawn from the post-replication pools of mixed (μ = 1.0), partially mixed (μ = 0.1), or unmixed (μ = 0.0) populations. Coinfections were performed with no mixing (μ = 0.0) or full mixing (μ = 1.0). **(B)** Ratios of weighted population fitnesses, compared between the second and the first round of infection. Second-round coinfections were initiated by viruses drawn from mixed, partially mixed, or unmixed populations, and were simulated under full mixing or unmixed conditions. **(C)** Overall proportions of parental and novel genotypes among the strains selected for second-generation replication (“pre-replication”), and among the populations resulting from a second generation of replication at μ = 0.0 (“post-replication”).

We observed a similar relationship when considering population fitness **(Figure 6B)**, and noted that, under unmixed conditions, the post-replication populations of viruses drawn from the μ = 0.1 pool had slightly higher fitnesses than the populations derived from the μ = 0.0 pool (median = 0.210 vs. median = 0.187; mean = 0.274 vs. mean = 0.257). These differences were not statistically significant — however, because the μ = 0.1 populations had experienced gene flow in their first generation of replication, we wanted to determine whether the slight gains in fitness and replication capacity were driven by selection on novel genotypes. We compared the proportion of novel genotypes in the starting populations (drawn from each first-generation replication pool) versus the resulting populations after a second round of infection under unmixed conditions **(Figure 6C)** and observed that selection in both the *μ* = 0.1 and the *μ* = 1.0 populations acted to increase the proportion of parental genotypes. Our findings reflect the observation that the generation of reassortant genotypes does not always lead to fitness increases in subsequent generations and suggest that reassortant-rich populations may frequently see reduced fitness compared to populations composed mainly of parental genotypes.

Finally, we repeated the next-generation simulations under the condition of epistasis between gene segments **(Supplemental Figure 3)**. In this scenario, the viruses drawn from the *μ* = 1.0 pools saw significant decreases in replication and fitness compared to those drawn from the *μ* = 0.0 pool, and the viruses drawn from the *μ* = 0.1 pool no longer showed any replication or fitness advantages compared to the *μ* = 0.0 pool. These results further demonstrate how inter-segment epistasis can lead to detrimental outcomes during multi-strain coinfections, and indicate that these negative fitness consequences may be especially pronounced during coinfections between viruses that are more distantly related.

## Discussion

We used a simulation approach to investigate how the processes of coinfection and reassortment can influence the fitness of segmented virus populations under different conditions. We observed that inter-strain mixing tended to have negative consequences for viral replication over a single generation across a variety of scenarios, with higher degrees of mixing corresponding to more substantial fitness costs. Conditions that exacerbated these costs included the existence of negative epistasis between gene segments and negative dominance dynamics between coinfecting alleles. Under certain conditions — for example, positive dominance dynamics, or in specific scenarios of immune pressure — we observed that mixing resulted in increased overall viral replication. However, outside of these specific circumstances, we found that inter-strain mixing generally resulted in both decreased replication and the persistence of low-fitness alleles.

While populations in the unmixed and mixed populations all contained the same number of initiating genomes, increased replication was achieved under unmixed conditions through the concentration of high-fitness genomes in one environment. Our finding that mixing between high- and low-fitness genomes causes an overall negative fitness effect reflects the findings of previous work describing how viral replication is subject to positive density dependence^64^ — positive density-dependent replication gains can be achieved by initiating infection with a greater number of virions, or by infecting with higher-fitness genotypes that will replicate more quickly, effectively increasing the intracellular multiplicity of infection. In both scenarios, increased replication is achieved through the structuring of resources (either high-fitness genomes or multiple identical viruses) in one area.

As expected, we also observed that inter-segment epistasis between viral genes led to greater negative consequences for heterologous coinfections. When strain mixing brings together interacting genes that are functionally mismatched, replication efficiency goes down and reassortant genotypes are more likely to have lower fitness. However, one limitation of this study is that we assume that the degree of negative epistasis correlates directly with the phylogenetic distance between mismatched strains. In nature, single point mutations may also impact the structure and function of proteins in a way that leads to negative inter-segment epistasis between closely-related strains^52^. Additionally, reassortment between distant strains may sometimes have minimal consequences: for example, the characterization of reassortant influenza viruses generated by crosses between avian H5N1 and human H3N2 indicated a high degree of compatibility between the two strains^27^. Contrastingly, reassortment between human H1N1 (which, similarly to H5N1, encodes a group 1 hemagglutinin and an N1 neuraminidase) and human H3N2 has been shown to generate low-fitness progeny^10^. These observations suggest that there may be intrinsic properties of certain viral genotypes that allow for reduced negative interactions between strains and indicate that further research is needed to characterize the role of inter-segment epistasis in reassortment for different viruses.

We also saw that negative dominance dynamics between alleles exacerbated the consequences of mixing between strains, which may have relevant implications for the behavior of non-standard viral genomes (nsVGs) during coinfections. nsVGs are observed across a wide range of RNA viruses, and have been shown to inhibit the replication of wild-type viruses in a dominant-negative manner^3,63,70–72^. nsVGs often contain large deletions that render key genes nonfunctional and are unable to replicate in the absence of a coinfecting helper virus — therefore, limiting coinfection between strains can also serve to curb the spread of these dominant-negative genotypes^72^.

In addition to nsVGs, segmented viruses (including influenza and bunyaviruses) may also produce semi-infectious virions that lack entire gene segments^8,9,73–76^. Similarly to nsVGs, these semi-infectious particles require complementation to carry out a productive infection. However, because semi-infectious virions carry an incomplete set of genes, rather than a complete genome encoding certain nonfunctional genes, the propagation of semi-infectious genomes across generations does not depend on the replication and packaging of a specific segment. The fitness effects of rescuing incomplete genomes through complementation during coinfection therefore likely depend on the extent to which the tendency to generate incomplete genomes is a strain-dependent property (as opposed to a consequence driven by random packaging error). If the generation of incomplete genomes is modulated by specific alleles, then limiting coinfection could help to remove those alleles from the population. Alternatively, if incomplete genome production is not controlled through simple, discrete viral genetic determinants, then coinfection could serve to rescue functional genotypes. For influenza virus, the generation of incomplete genomes appears to be at least partially genetically encoded^3,9,51,77^. suggesting that viral populations may benefit from limiting interactions with genotypes that are more likely to generate incomplete genomes.

When we applied our model to a situation in which certain viral genes are subject to immune pressure, we found that the response to strain mixing depended both on the degree of immune resistance in the population and the antigenic structure of the virus. Under most of the modeled conditions, mixing had a negative or negligible impact on the replication and fitness of populations under immune pressure. However, we observed that when population resistance to immunity was at 15%, and both segments A and B were subjected to immune pressure, viral replication responded positively to increased strain mixing. The prevalence of standing immune resistant variants in the population is an important factor in determining the impact of mixing on fitness, because the fitness benefits of mixing are driven by the transfer of resistance alleles to non-resistant genotypes. At a low level of resistance, there are too few resistant alleles to make a major impact on the population, and at a high level of resistance, there are too few non-resistant genotypes that would benefit from accepting new resistance alleles. In keeping with this logic, we observed that there was only a small window of population resistance at which mixing was actively beneficial.

The fact that we observed a positive relationship between mixing and replication for AB < C but not C < AB viruses can be explained in part by the different reassortment outcomes that are possible under each condition. Under a C < AB antigenic scheme, when a single coinfecting partner carries a resistance allele, reassortment can result in the transfer of the allele to either a higher-fitness or a lower-fitness genome (depending on the fitness of the non-resistant coinfection partner). This reshuffling often has fitness consequences — for example, recombination can break positive associations between antibody escape mutations and mutations conferring increased transmissibility^78^, thereby redistributing beneficial variants across multiple separate genotypes. These same outcomes are possible under an AB < C scheme; however, when each coinfecting virus encodes a single resistance allele for a separate antigen, reassortment can also allow for the consolidation of these alleles on a single genome. Therefore, coinfecting viruses that are essentially inviable on an individual basis (because they each only carry one of the two necessary resistance alleles) can combine through reassortment to generate viable offspring. This outcome stands in contrast to the single-antigen scenario, where reassortment can cause viability to be spread to new genomes, but not generated *de novo* from two nonviable parental genomes.

Along with the immediate consequences for replication, we also observed significant differences in the allelic composition of populations whose strains were allowed to mix. Heterologous mixing relaxed the force of selection on lower-fitness mutations, allowing these variants to persist at higher frequencies than in the unmixed populations. This effect has been described previously in studies that examine how inter-strain coinfection can impact viral population diversity by supporting the persistence of minor variants^56^, contributing to population robustness^79^, and promoting the fixation of neutral alleles^80^. While these effects may have negative consequences during short-term evolution, the fitness effect of a given allele is frequently context-dependent, and a higher population diversity may serve as an advantage in more variable environments.

Additionally, despite the deleterious effects of increased mixing on population-level fitness, we observed that reassortment occasionally yielded novel individual genotypes with higher fitness than either parental strain. Coinfecting viruses were still able to generate some of these genotypes under a low mixing rate and were thereby able to access expanded genomic diversity while minimizing replication defects. However, in following generations of infection, viruses generated under these low-mixing conditions saw similar fitness outcomes as viruses generated under unmixed conditions, and positive selection acted primarily on parental genotypes. These results suggest that novel reassortant genotypes only occasionally promote sustained fitness benefits. In general, the conditions under which reassortment can generate novel, high-fitness genotypes are limited. If both parental strains carry low-fitness alleles for the same gene, then, in the absence of mutation, any reassortant genotype will contain a low-fitness variant of the gene in question. If one parent carries a high-fitness allele for the gene, it can be passed to the other strain — however, coinfection can support the replication of the lower-fitness allele through phenotypic hiding, rendering selection against the allele less efficient, and increasing the chances that it will persist regardless. Therefore, reassortment poses the greatest benefits when parental strains carry lower-fitness alleles on different segments, which they can then reciprocally swap to generate a genotype with higher fitness than either original strain.

Taken as a whole, our findings highlight the need for successful viruses to strike a balance between diversification and maximizing fitness in a single environment. One feature of many viral life cycles that could help to achieve this balance is bottlenecking during transmission, and both theoretical and experimental studies have shown how bottlenecks can promote efficient selection on standing viral diversity^81–83^. Another strategy (which may also serve to decrease bottleneck size) is superinfection exclusion. Previous work has described how superinfection exclusion can be beneficial on short timescales, but eventually leads to fitness consequences arising from decreased population adaptability^80^. However, depending on the viral system in question, superinfection exclusion may not result in the absolute inhibition of all secondary infections. For example, across influenza viruses, superinfection exclusion can result in varying degrees of inhibition depending on the relative timing of the two infections^37,38^ and the replication kinetics of the primary virus^39^. Therefore, superinfection exclusion may help minimize the negative consequences of unrestricted inter-strain mixing, while still allowing diversity to be generated and maintained by the smaller subset of cells that do experience successful coinfection. Successful viruses may explore different degrees and mechanisms of restriction to achieve an optimal compromise between diversification and selection.

## Methods

### Replication simulations

Replication events were simulated using a Gillespie algorithm^62^. Briefly, replication events were initiated at time intervals drawn from an exponential distribution with a mean of 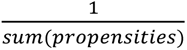, and the specific reaction to take place was randomly selected according to each reaction’s relative propensity. Simulations were performed for 8 time units, across each of the 100 genotype pairs. As the time steps are defined in arbitrary units, the number of steps was selected so that the final number of replicated genomes per simulation roughly aligned with estimates for the number of influenza genomes generated during a single cycle of infection^84^. Simulations were performed with optimized tau-leaping (*ε* = 0.01) in *R (v4.1.1)* using the *GillespieSSA* package^85^.

### Packaging simulations

Gillespie simulations were performed using the *GillespieSSA* package for 1000 time steps across all 100 coinfecting pairs. The packaging propensities for each possible genotype were defined as:

**Table 2.** Per-genotype packaging propensities.

| Packaging reaction | Propensity |
| --- | --- |
| $A_i B_i C_i \rightarrow A_i B_i C_i + 1$<br>$A_i \rightarrow A_i - 1$<br>$B_i \rightarrow B_i - 1$<br>$C_i \rightarrow C_i - 1$ | $p_{A_i} * p_{B_i} * p_{C_i}$ |
| $A_i B_i C_j \rightarrow A_i B_i C_j + 1$<br>$A_i \rightarrow A_i - 1$<br>$B_i \rightarrow B_i - 1$<br>$C_j \rightarrow C_j - 1$ | $p_{A_i} * p_{B_i} * p_{C_j}$ |
| $A_i B_j C_i \rightarrow A_i B_j C_i + 1$<br>$A_i \rightarrow A_i - 1$<br>$B_j \rightarrow B_j - 1$<br>$C_i \rightarrow C_i - 1$ | $p_{A_i} * p_{B_j} * p_{C_i}$ |
| $A_i B_j C_j \rightarrow A_i B_j C_j + 1$<br>$A_i \rightarrow A_i - 1$<br>$B_j \rightarrow B_j - 1$<br>$C_j \rightarrow C_j - 1$ | $p_{A_i} * p_{B_j} * p_{C_j}$ |
| $A_j B_i C_i \rightarrow A_j B_i C_i + 1$<br>$A_j \rightarrow A_j - 1$<br>$B_i \rightarrow B_i - 1$<br>$C_i \rightarrow C_i - 1$ | $p_{A_j} * p_{B_i} * p_{C_i}$ |
| $A_j B_i C_j \rightarrow A_j B_i C_j + 1$<br>$A_j \rightarrow A_j - 1$<br>$B_i \rightarrow B_i - 1$<br>$C_j \rightarrow C_j - 1$ | $p_{A_j} * p_{B_i} * p_{C_j}$ |
| $A_j B_j C_i \rightarrow A_j B_j C_i + 1$<br>$A_j \rightarrow A_j - 1$<br>$B_j \rightarrow B_j - 1$<br>$C_i \rightarrow C_i - 1$ | $p_{A_j} * p_{B_j} * p_{C_i}$ |
| $A_j B_j C_j \rightarrow A_j B_j C_j + 1$<br>$A_j \rightarrow A_j - 1$<br>$B_j \rightarrow B_j - 1$ | $p_{A_j} * p_{B_j} * p_{C_j}$ |
| $C_j \rightarrow C_j - 1$ | |

## Acknowledgments

We are grateful to Dr. James O’Dwyer for guidance in the initial development of the study and Dr. Pamela Martinez for critical reading of manuscript drafts. C.B.B was supported by NIH R01AI139246, NIH R01AI179910, and 1U01AI186993. M.F. was supported in part by NIH U19AI181767.

## Data availability

Code and data involved in this project can be found at https://github.com/BROOKELAB/model-coinfection.

## Supplemental Figures

**Supplemental Figure 1.**
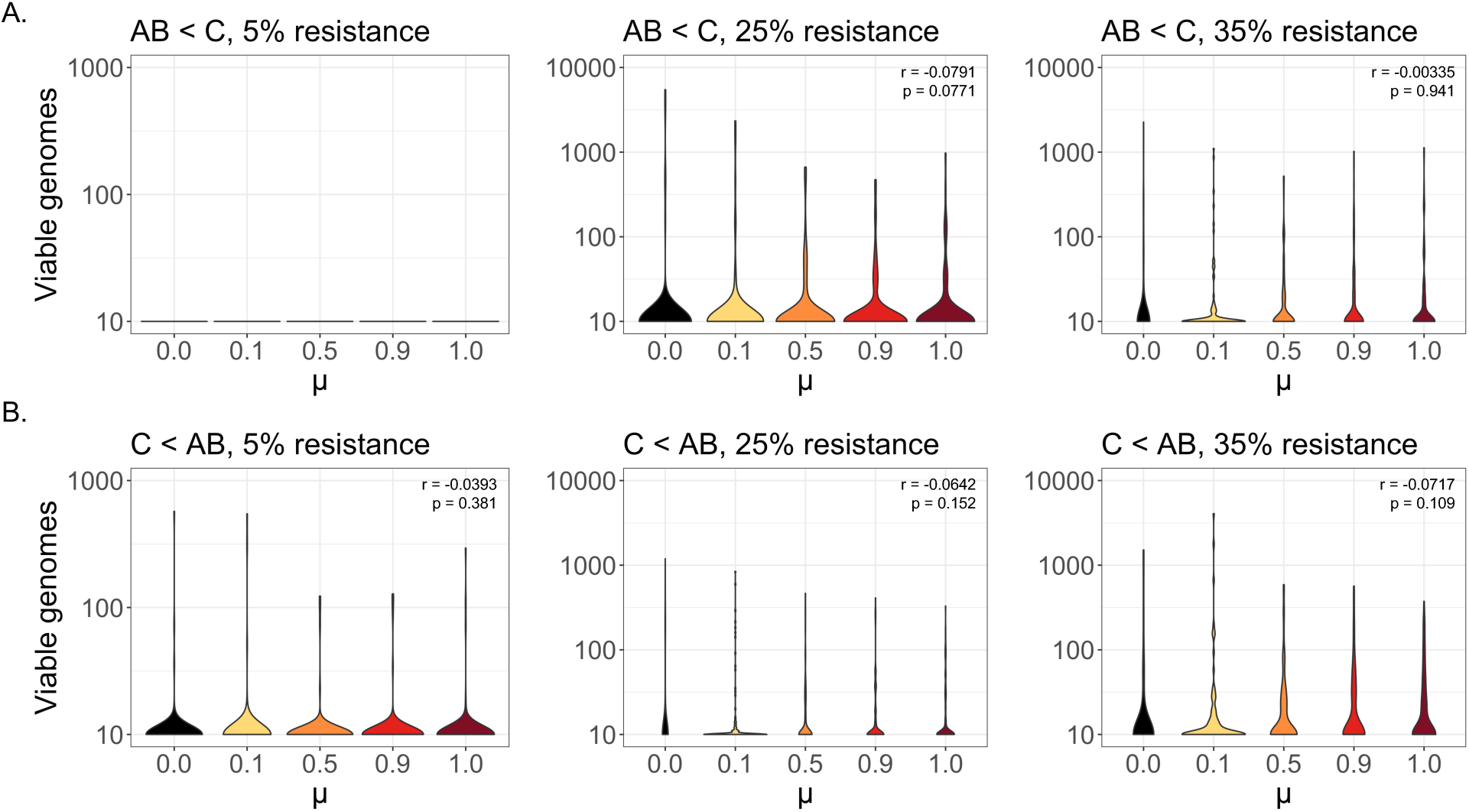
The impact of mixing on replication under different antigenic schemes and different levels of population immunity. **(A)** AB < C viable genome counts for increasing values of μ, under varying population resistance levels. **(B)** C < AB viable genome counts for increasing values of μ, under varying population resistance levels.

**Supplemental Figure 2.**
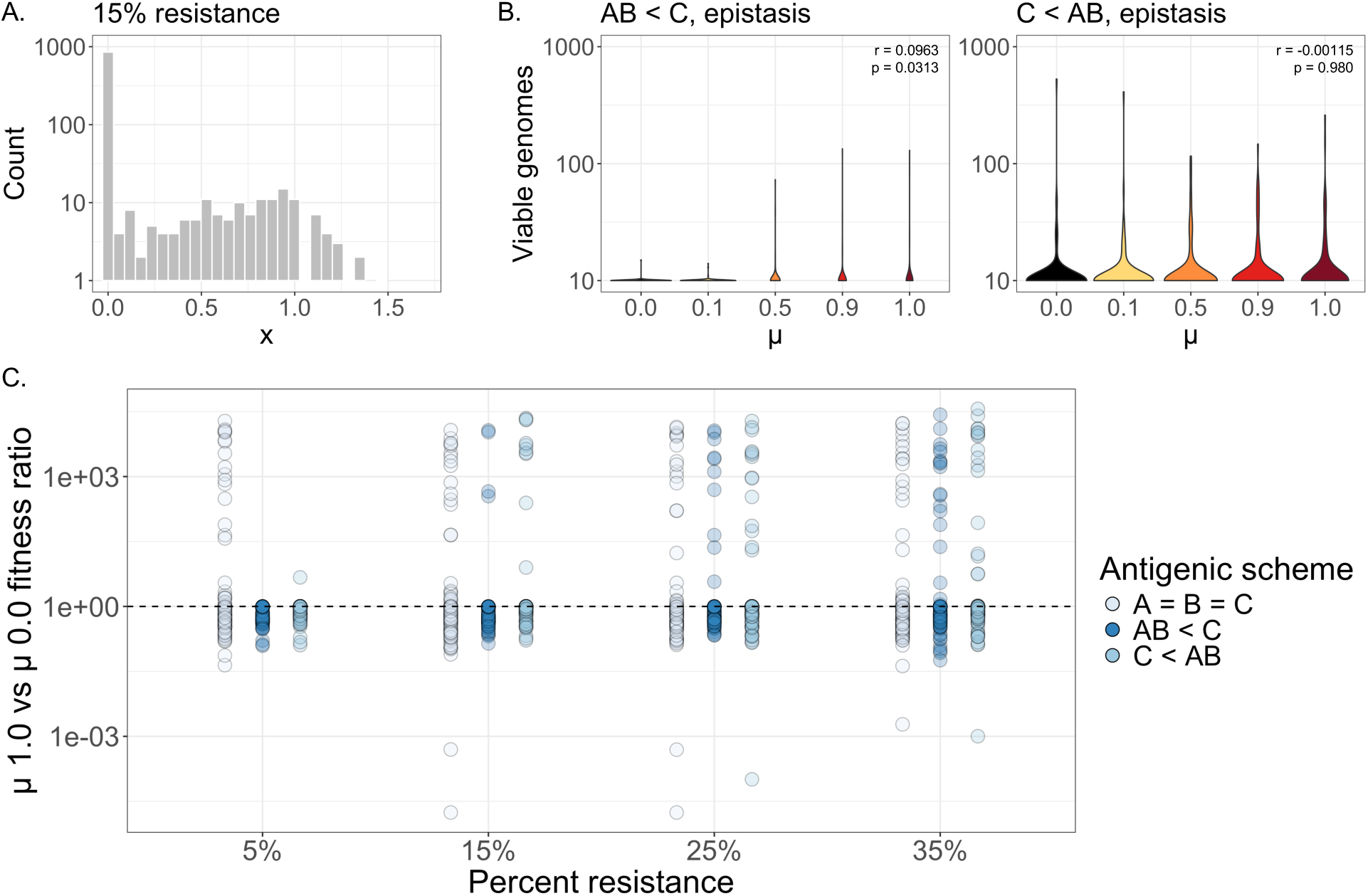
The impact of immune pressure on replication and fitness under conditions of inter-segment epistasis. **(A)** Distribution of gene activity (x) values under an immune-pressure model, where 15% of variants encode some degree of resistance. **(B)** Total viable genomes under different mixing rates (μ), for AB < C and C < AB antigenic schemes, under epistatic conditions. Insets show *ρ* and p-values, derived from a Spearman’s rank correlation test. **(C)** Ratio of weighted population fitness scores between fully mixed (μ = 1.0) and unmixed (μ = 0.0) populations, under epistatic conditions, shown in the absence of immune pressure (A = B = C) or under different immunity schemes (AB < C or C < AB) under varying levels of population resistance. Dashed line represents a fitness ratio of 1.0.

**Supplemental Figure 3.**
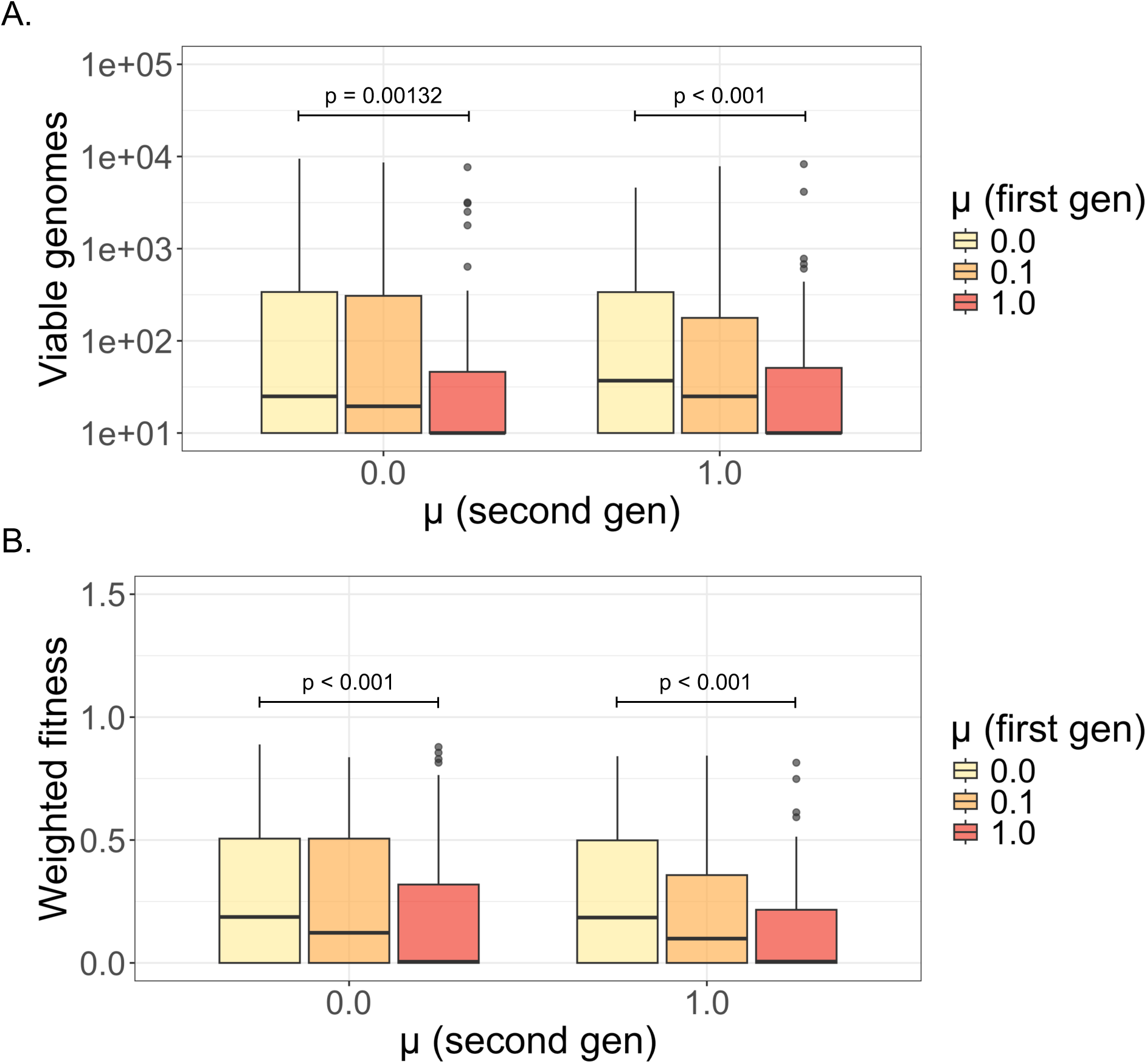
The impact of mixing in a second generation of replication, under conditions of inter-segment epistasis. **(A)** Total viable genomes from coinfections initiated by viruses drawn from the post-replication pools of mixed (μ = 1.0), partially mixed (μ = 0.1), or unmixed (μ = 0.0) populations (under epistatic conditions). Coinfections were performed with no mixing (μ = 0.0) or full mixing (μ = 1.0). P-values are derived from Mann-Whitney U tests. **(B)** Ratios of weighted population fitnesses, compared between the second and the first round of infection, under epistatic conditions. Second-round coinfections were initiated by viruses drawn from mixed, partially mixed, or unmixed populations, and were simulated under full mixing or unmixed conditions. P-values are derived from Mann-Whitney U tests.

